# Resilience of multispecies biofilm communities: Comparative analysis of a *Veillonella dispar*-dominant oral biofilm under diverse oxygen conditions *in vitro*

**DOI:** 10.64898/2026.09.25.754398

**Authors:** Yue Sun, Teresa Lea Nguyen, Katharina Doll-Nikutta, Nils Heine, Guntram A. Grassl, Carina Mikolai, Andreas Winkel, Meike Stiesch

## Abstract

Oral multispecies biofilms developing on dental implants drive peri-implant mucositis and periimplantitis. They are shaped by the physicochemical conditions of their surrounding environment, among which oxygen availability is an important influential factor. However, while oxygen tension fluctuates significantly within the peri-implant pocket, whether these environmental shifts are sufficient to influence the structure and composition of an established microbial community remains unclear. For this purpose, we evaluated the structure and composition of a four-species oral biofilm model comprising the species *Streptococcus oralis*, *Actinomyces naeslundii*, *Veillonella dispar* and *Porphyromonas gingivalis*, which was cultivated under normoxic (21% O□), hypoxic (1% O_2_) and anoxic (0% O□) conditions for a period of 21 days. Biofilm 3D architecture, volume and membrane integrity was analyzed by LIVE/DEAD^®^ staining combined with confocal laser scanning microscopy. Community composition was assessed by quantitative real-time PCR and complemented by Fluorescence *In Situ* Hybridization (FISH) to provide structural context. While normoxia transiently promoted higher early biofilm volume and membrane integrity, the community exhibited remarkable long-term resilience, consistently maintaining a *V. dispar*-dominated structure across all oxygen gradients. Crucially, the obligate anaerobic pathogen *P. gingivalis* persisted as a viable minority (∼1%) even under 21-day normoxic cultivation, sheltered within the deeper layers of the biofilm. These results suggest that clinical oxygen levels alone are not able to drastically modulate established oral biofilm communities and that oxygen-sensitive pathogens persist in a subclinical sanctuary protected by the surrounding non-pathogenic species.

## Introduction

Dental implants have become a cornerstone of modern oral rehabilitation, providing predictable restoration of function and aesthetics after tooth loss (Pye, 2009). While the long-term success rates of dental implants are generally high (Quirynen, 2002), peri-implant diseases such as peri-implant mucositis and peri-implantitis remain common and clinically significant complications that can reduce implant longevity (Zitzmann, 2008) (Mombelli A. M., 2012) (Pott, 2025). These diseases are driven by biofilm-mediated infections (Mombelli A. V., 1987) (Schwarz, 2018), in which complex microbial communities are embedded in an extracellular matrix that enhances tolerance to environmental stress, antimicrobial agents, and host immune defences (Flemming H. C., 2016) (González, 2018).

Biofilms developing on implants are shaped by the physicochemical conditions of their surrounding environment, among which oxygen availability is an important influential factor (Knighton, 1984) (Antoniazzi, 2016). Under physiological conditions, different tissues are characterized by distinct oxygen partial pressures that reflect their metabolic activity, a state commonly referred to as physiological normoxia or physioxia (Carreau, 2011) (Mirchandani, 2025). While supragingival surfaces are typically exposed to atmospheric oxygen, the peri-implant sulcus becomes hypoxic as a result of cell oxygen consumption and restricted diffusion. Thorn *et al*. reported a transmucosal oxygen concentration of 5.3–5.5% (39.8–41.2 mmHg) at the surface of oral tissues (Thorn, 1997), and Mettraux *et al*. reported an average oxygen tension of 1.8% O_2_ (13.3 mmHg) within untreated periodontal pockets (Mettraux G. R., 1984). During infection, the proliferation of pathogens combined with immune cell infiltration leads to a marked reduction in local oxygen levels, which can drop to as low as 0.1% (Taylor CT, 2007). These oxygen fluctuations are not merely passive background conditions, but rather represent a selective pressure thought to influence microbial physiology, spatial organization, and pathogenic potential. Ultimately, this pressure dictates whether a commensal community can resist the encroachment of anaerobic pathogens or if the environmental shift facilitates a transition to a disease-associated state (Loesche, 1983) (Takahashi, 2015) (Valm, 2019).

Although oxygen tension is known to influence microbial succession and gene expression, few *in vitro* studies have systematically investigated its long-term effects on multispecies biofilm formation. Most established models operate under aerobic or fully anaerobic conditions, overlooking the physiologically relevant hypoxic states common in peri-implant sites. Hypoxia, in particular, represents a transitional and potentially stable state in diseased tissues, yet remains underexplored in oral biofilm research.

In this study, we therefore investigated the structure and composition of a four-species oral biofilm model composed of *Veillonella dispar*, *Actinomyces naeslundii*, *Streptococcus oralis*, and *Porphyromonas gingivalis*, under three oxygen conditions: normoxic (21% O□), hypoxic (1% O□), and anoxic (0% O□). The biofilms were cultivated on the implant material titanium and glass control surfaces for 21 days, enabling the analysis of both early and mature stages of community development. Using LIVE/DEAD^®^ staining and Fluorescence *In Situ* Hybridization (FISH) followed by confocal laser scanning microscopy (CLSM) as well as quantitative real-time PCR (qRT-PCR), we characterized biofilm volume, viability, and species composition across time and oxygen conditions. By dissecting the influence of oxygen availability on biofilm behavior, this work provides insight into the ecological and structural adaptations of mature oral microbial communities under shifting environmental pressures and support deciphering the early process of infection development.

## Materials and Methods

### Bacteria strains and pre-culture

*Streptococcus oralis* ATCC^®^ 9811™, *Actinomyces naeslundii* DSM 43013, *Veillonella dispar* DSM 20735 and *Porphyromonas gingivalis* DSM 20709 were cultivated on fastidious anaerobe agar (FAA; Neogen Corporation, Lansing, Michigan, USA) plates with sheep blood (Thermo Fisher, Waltham, Massachusetts, USA) for 72 h. The colonies were then separately precultured anaerobically for 24 h in brain heart infusion broth (BHI; Oxoid, Wesel, Germany), supplemented with 10 µg/mL vitamin K1 (Roth, Karlsruhe, Germany) and 5 mg/L hemin (Sigma-Aldrich, Burlington, Massachusetts, USA) at 37 °C.

### Co-culture and biofilm formation

The precultures were diluted in fresh BHI + vitamin K1 + hemin medium to an initial optical density (OD_600_) of 0.1. Equal volumes were combined, further diluted to OD_600_ = 0.01, which corresponds to 1.17×10^7^ CFU/mL *S. oralis*, 1.63×10^6^ CFU/mL *A. naeslundii*, 3.24×10^5^ CFU/mL *V. dispar* and 7.38×10^4^ CFU/mL *P. gingivalis* (Kommerein, 2017). 1 mL aliquots were applied to Ø 12 mm titanium or glass discs placed in 24-well plates (Greiner, Frickenhausen, Germany). Titanium was selected as a clinically relevant implant substrate, while glass was utilized in parallel to minimize metallic reflection and optimize optical resolution during fluorescence microscopy. Incubation was performed at 37 °C under normoxic (21% O□, 5% CO_2_), hypoxic (1% O_2_, 5% CO_2_, 94% N_2_) and anoxic (0% O□, 5% CO_2_) conditions. Biofilms were evaluated at multiple time points (1, 2, 7, 14 and 21 days). For each 2-3 days during the incubation, 0.5 mL of the mixed bacterial culture was replaced with 0.6 mL of fresh medium to simulate *in vivo* exchange of nutrition and compensate for evaporation-induced volume loss.

### LIVE/DEAD^®^ staining and confocal laser scanning microscopy analysis

Biofilms were harvested at 1, 2, 7, 14 and 21 days of incubation in 6-well plates and stained using SYTO^®^9 and propidium iodide (LIVE/DEAD^®^ BacLight™ Bacterial Viability Kit, Thermo Fisher, Waltham, Massachusetts, USA) at final concentrations of 1.67 µM and 10 µM, respectively.

Stained biofilms were covered in PBS and visualized using a confocal laser scanning microscope (CLSM) (SP-8, Leica Microsystems, Wetzlar, Germany) with 63× objective. Detection of SYTO^®^9 fluorescence was performed using a 488 nm excitation wavelength from a laser, with emission captured between 500–540 nm. Propidium iodide was excited at 552 nm, and its emission was recorded within the 675–750 nm range. Confocal image stacks were acquired at 1024 × 1024-pixel resolution, spanning from bottom to top of each biofilm. Five regions were imaged from each biofilm. The biofilm surface was reconstructed, and the enclosed volume was quantified using Imaris software (version 8.4, Bitplane, Zurich, Switzerland).

### Fluorescence *In Situ* Hybridization

Biofilms were fixed with 50% ice-cold ethanol and air-dried. Samples were then treated with 100 μL lysozyme solution (1□µg/µL) at 37□°C to permeabilize Gram-positive bacteria. Following enzymatic treatment, ethanol washes (2 × 100%) were applied to stop lysis, and samples were air-dried.

Hybridization was performed using staining solution containing urea-NaCl buffer (1 M urea, 0.9 M NaCl, 20 μM Tris-HCl, pH 7.0) and fluorescently labeled FISH probes (Table S-1). The samples were incubated with the staining solution at 46 °C for 30 minutes in a moist chamber. After hybridization, samples were washed with wash buffer (4 M urea, 0.9 M NaCl, 20 μM Tris-HCl, pH 7.0) at 48□°C thrice and rinsed with distilled water. For imaging, biofilms were covered with PBS and visualized using confocal laser scanning microscopy (CLSM) (SP-8, Leica Microsystems, Wetzlar, Germany), in which fluorescence signals were acquired in two sequences. In the first, ALEXA Fluor^®^ 405 and 568 signals were detected using 405 nm and 552 nm lasers with emission ranges of 413–477 nm and 576– 648 nm, respectively. In the second, ALEXA Fluor^®^ 488 and 647 signals were activated with 488 nm and 638 nm lasers (emission: 509–576 nm and 648–777 nm, PMT detectors). Image Z-stacks were recorded at 2 μm intervals and processed using Imaris software (version 8.4, Bitplane, Zurich, Switzerland). Confocal image stacks were acquired at 1024 × 1024-pixel resolution, spanning from bottom to top of each biofilm. Five regions were imaged from each biofilm.

### Propidium monoazide (PMA) treatment and DNA isolation

To quantify compositions of viable parts of each bacteria species inside the biofilms, the biomass was collected and treated with PMA, and analyzed using quantitative real time polymerase chain reaction (qRT-PCR). PMA selectively penetrates cells with compromised membrane integrity, where it binds to DNA and inhibits subsequent PCR amplification (Àlvarez, 2013). In contrast, it is excluded from viable cells with intact membranes. Therefore, only viable cells are detected during qRT-PCR.

Multispecies biofilms were harvested by scraping with a sterile cell scraper after washing with PBS. The detached biomass was collected in 2 mL microcentrifuge tubes, washed twice with PBS at 4□°C, and resuspended in 200□µL PBS. 50□µL of the suspension was transferred to 1.5□mL tubes. PMAxx (2□mM) (Biotum, Hayward, California, USA) was freshly prepared in PCR-grade water and homogenized. A volume of 6 µL PMAxx (final concentration 240 µM) was added to each sample and mixed thoroughly. Samples were incubated for 10 min at 4□°C in the dark, followed by 20 min exposure to 470 nm blue LED light. Bacteria were then washed with PBS to remove unbound PMA.

The bacterial genomic DNA was extracted using the QIAamp DNA Mini Kit (Qiagen, Hilden, Germany) with mechanical and enzymatic lysis. PMA treated biofilm samples were centrifuged and resuspended in 450□µL lysozyme buffer (20□mg/mL lysozyme, 20□mM Tris-HCl, pH 8.0, 2□mM EDTA, 1.2% Triton X-100). Samples were incubated at 37□°C for 1–2 hours, followed by the addition of 50□µL proteinase K and 500□µL AL buffer. After mixing, samples were incubated at 56□°C for 30 min and then at 95□°C for 15 min. Lysates were cooled on ice for 5 min. Afterwards, the samples were then transferred to Lysing Matrix E tubes (MP Biomedicals, Santa Ana, California, USA) and subjected to bead milling using a Precellys 24 homogenizer at 6500 rpm for 30 s, repeated three times with cooling intervals. Supernatants were then processed following the instructions of Qiagen. Purified DNA was eluted in 50 µL PCR-grade water, quantified using a Nanodrop 2000c photometer (Thermo Fisher, Waltham, Massachusetts, USA) and stored at −20 °C until further analysis.

### Quantitative real-time PCR

Quantitative real-time PCR (qRT-PCR) was used to assess the relative abundance of the four species within the multispecies biofilms, using a Roche Lightcycler 96 (Roche, Basel, Switzerland). Same species-specific primers were used as described before (Kommerein, 2017). A master mix of each reaction contained 12.5 μL iQ™ SYBR^®^ Green Supermix solution (BioRad, Hercules, California, USA), 0.2 μM forward and reverse primer and 1-40 ng of template DNA. The reactions were run with an initial denaturation at 95□°C for 3 min, followed by 40 cycles of denaturation at 95□°C for 10 s, species-specific annealing (58□°C for *S. oralis*, *A. naeslundii*, and *V. dispar*; 56□°C for *P. gingivalis*) for 20 s, and extension at 72□°C for 20 s. Following amplification, a melting curve analysis was performed at 60□°C for 6□s over 115 cycles. For each bacterial species, a standard curve was established using serial dilutions of genomic DNA at defined concentrations. All qRT-PCR reactions were performed in technical triplicate. The amount of genomic DNA for each target species in the unknown samples was calculated from the corresponding standard curve. To estimate the number of bacterial cells, the measured DNA quantity was divided by the genome weight per cell for the respective species.

### Statistical analysis

Statistical analysis was performed using GraphPad Prism software version 10.6.1 (GraphPad Software, Boston, Massachusetts, USA). Data were analyzed using a two-way analysis of variance (ANOVA) to evaluate the primary and interactive effects of cultivation time and oxygen condition on biofilm parameters, followed by a Tukey’s post-hoc test for multiple comparisons and p-value correction, with statistical significance defined at α = 0.05.

## Results

### Early-stage biofilm volume is transiently influenced by oxygen availability

Multispecies biofilm consisting of *S. oralis*, *A. naeslundii*, *V. dispar* and *P. gingivalis* were cultivated on Ø 12 mm glass or titanium discs in 24-well plates under anoxic (0%), hypoxic (1%), and normoxic (21%) conditions, and collected at days 1, 2, 7, 14, and 21. The collected biofilms were stained with the LIVE/DEAD^®^ viability kit and immediately analyzed by CLSM. The CLSM images of the biofilms are depicted in Fig. 1 as 2D maximum intensity projections and side views from the 3D reconstruction.

**Fig. 1.**
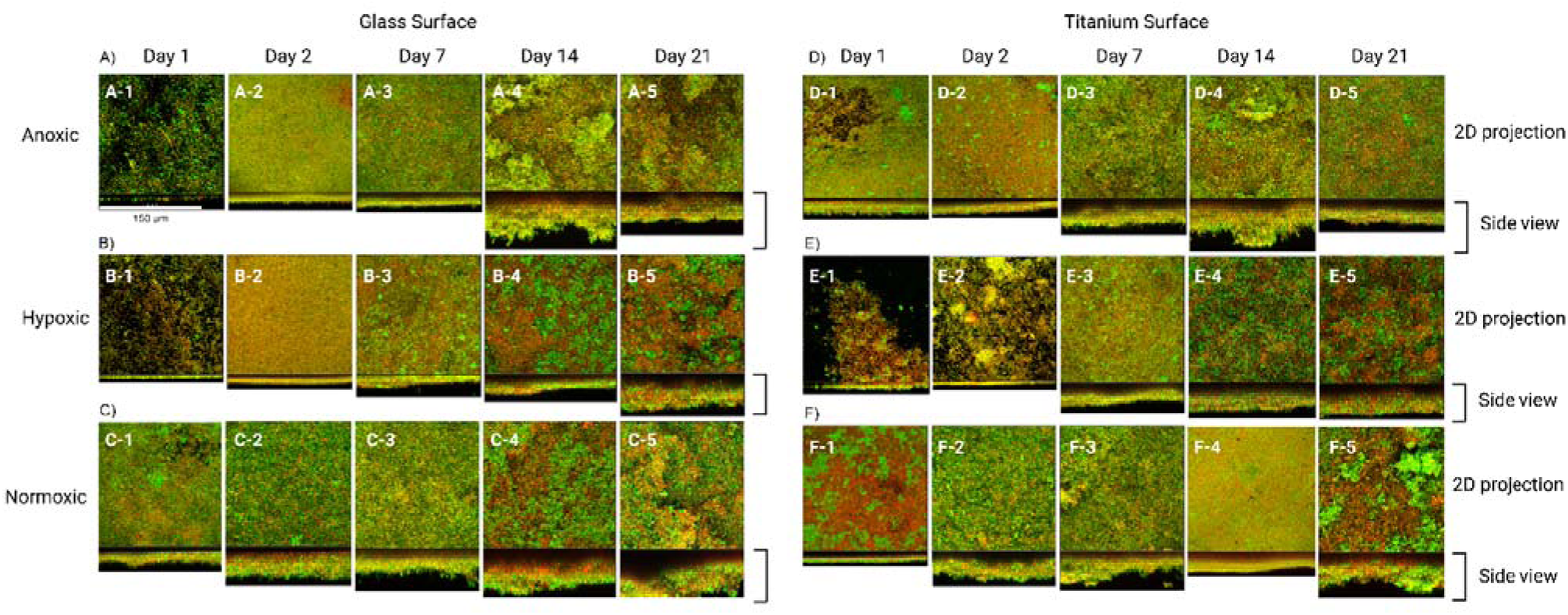
Representative microscopy images of LIVE/DEAD^®^-stained biofilms cultivated for 1, 2, 7, 14 and 21 days on A) - C) glass or D) - F) titanium surfaces under A) D) anoxic (0%), B) E) hypoxic (1%), C) F) normoxic (21%) conditions. Viable bacteria with intact membranes are visualized in green and membrane-compromised bacteria are visualized in orange/red. Scale bar = 150 μm.

Biofilm thickness increased progressively from day 1 to day 7, reaching its maximum between days 14 and 21. While biofilms cultivated on glass substrates developed a heterogeneous topography with high-viability, mound-like structures and localized clusters starting on day 7, titanium surfaces displayed this irregular, clustered morphology as early as day 1.

The biofilm volumes from these images were calculated using the Imaris software and are presented in Fig. 2. Biofilm volume increased from day 1 to 7 under all oxygen conditions on both surfaces, and reached its peak between day 7 and 21. On glass surfaces, no statistically significant differences in biofilm volume were observed between oxygen conditions at day 1 (Fig. 2 A). By day 2, normoxic biofilms on glass exhibited significantly greater volumes than those cultivated under anoxic (approx. two-fold, p = 0.0114) and hypoxic (approx. two-fold, p = 0.0002) conditions. At day 7, biofilm volumes increased across all oxygen conditions, and the volume differences between the conditions were not notable. At day 14, the biofilm volumes plateaued for all conditions, suggesting a shift in microbial density and reduced growth dynamics. Under these conditions, the anoxic biofilm on glass exhibited a higher volume compared with the hypoxic biofilm. By day 21, no marked differences in overall biofilm volume were observed among the oxygen conditions.

**Fig. 2.**
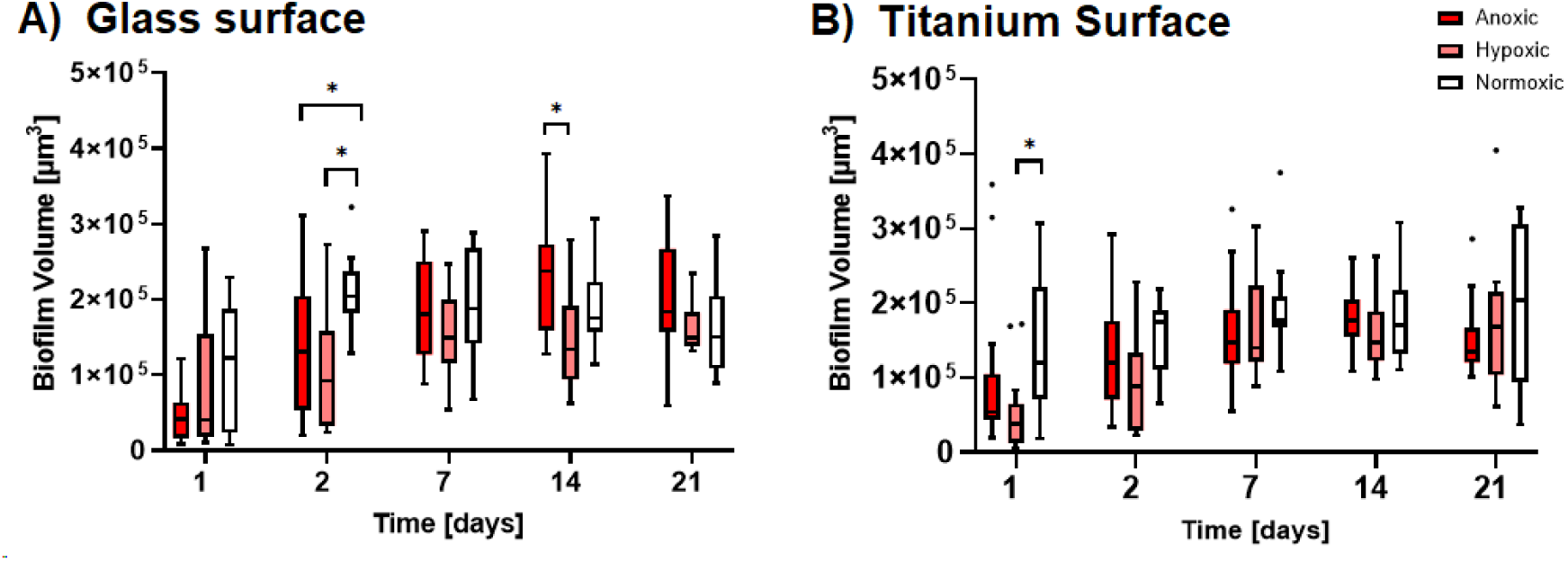
Tukey box plots of biofilm volume on A) glass and B) titanium surfaces under anoxic, hypoxic, and normoxic conditions over a 21-day cultivation period. * indicates statistically significant differences with p ≤ 0.05.

On titanium surfaces, normoxic conditions resulted in significantly higher biofilm volumes than hypoxic conditions at day 1 (approx. three-fold, p = 0.0304) (Fig. 2 B). By day 2, however, the differences between oxygen conditions became less notable, and no statistically significant variations were detected. As observed on glass surfaces, biofilm volumes on titanium surface also increased across all conditions by day 7. Another similar trend to that on glass was observed at day 14: the overall volumes plateaued across conditions, suggesting reduced growth dynamics. Likewise, by day 21 no marked differences in biofilm volume were detected among the oxygen environments.

Overall, biofilm volume differed between oxygen conditions during the early stages (days 1–2) of development with increased biofilm growth under normoxic conditions, whereas these differences were rarely apparent after day 7.

### Biofilm membrane integrity differs between normoxic and oxygen-limited conditions

Besides biofilm morphology and volume, LIVE/DEAD® staining could also be used to analyze bacterial membrane integrity as parameter of cellular viability, revealing differences associated with oxygen conditions and surface type (Fig. 3). On glass surfaces, biofilms cultivated under anoxic conditions exhibited significantly higher membrane integrity at day 1 (approx. 68 %) compared with hypoxic conditions (approx. 45%, p = 0.0092), but no significant difference were observed relative to normoxic conditions. From day 2 onwards, normoxic biofilms exhibited consistently higher membrane integrity (approx. 70-75%, Fig. 3A). This difference was most substantial on day 2, significantly exceeding both anoxic (p = 0.0003) and hypoxic (p = 0.0006) conditions (Tab. 1). On day 7, the gap between biofilms under various conditions narrowed. Nevertheless, normoxic biofilms still showed a higher membrane integrity compared to hypoxic biofilms with a p-value of 0.0403. At days 14 and 21, membrane integrity again diverged more clearly between normoxic and oxygen-limited conditions. Overall, with the exception of day 1, biofilms consistently followed a similar pattern throughout the cultivation period: normoxic conditions were associated with higher proportions of cells with intact membranes, whereas hypoxic and anoxic conditions showed increased proportions of membrane-compromised cells.

**Fig. 3.**
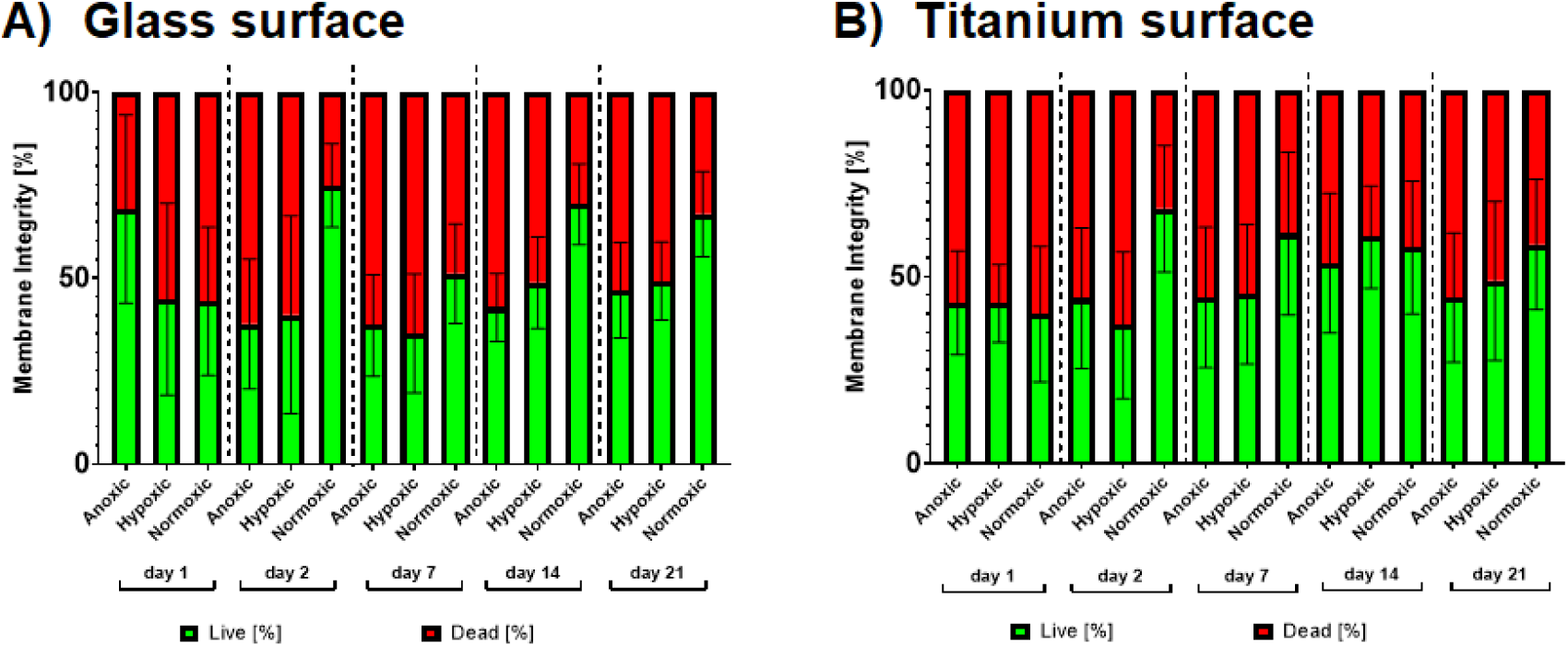
Mean and standard deviation of biofilm membrane integrity on A) glass and B) titanium surfaces under anoxic, hypoxic, and normoxic conditions over a 21-day cultivation period. Statistical results can be found in Tab. 1.

**Table 1:** 2way ANOVA comparisons of biofilm membrane integrity between different oxygen conditions, followed by a Tukey’s post-hoc test for multiple comparisons and p-value correction, with statistical significant differences defined at α = 0.05. For the complete statistical results see table S-2 in the supplementary section

|  |  | Comparison | P |
| --- | --- | --- | --- |
| On glass surface |  |  |  |
|  | Day 1 | Anoxic vs. hypoxic | 0.0092 |
|  | Day 2 | Anoxic vs. normoxic | 0.0003 |
|  |  | Hypoxic vs. normoxic | 0.0006 |
|  | Day 7 | Hypoxic vs. normoxic | 0.0403 |
|  | Day 14 | Anoxic vs. normoxic | < 0.0001 |
|  |  | Hypoxic vs. normoxic | < 0.0001 |
|  | Day 21 | Anoxic vs. normoxic | 0.0219 |
|  |  | Hypoxic vs. normoxic | 0.0091 |
| On titanium surface |  |  |  |
|  | Day 2 | Anoxic vs. normoxic | 0.0042 |
|  |  | Hypoxic vs. normoxic | 0.0008 |

Biofilms formed on titanium surfaces exhibited overall trends comparable to those on glass, generally showing higher membrane integrity under normoxic conditions (approx. 60-65%, Fig. 3B). However, statistical significance across time points was less consistent, with notable exceptions occurring on days 1 and 14 (Tab. 1). No significant differences between surface types were detected at these time points. At day 2, however, normoxic biofilms on titanium also demonstrated significantly higher membrane integrity compared with both anoxic and hypoxic conditions (both approx. 45%, p = 0.0042 and p = 0.0008, respectively).

Overall, membrane integrity differed primarily between normoxic and oxygen-limited environments, whereas hypoxic and anoxic biofilms exhibited largely comparable profiles throughout the cultivation period.

### *V. dispar* remains the dominant species throughout biofilm development across oxygen conditions

Quantitatve RT-PCR was performed to determine the relative proportions of each species within the four-species biofilms cultivated on glass surfaces over a 21-day period under anoxic, hypoxic, and normoxic conditions (Fig. 4). Based on the previous comparable results and as preliminary qRT-PCR analyses of biofilms cultivated on titanium surfaces during the early stages (days 1–2; one biological replicate) revealed species distributions comparable to those observed on glass surfaces, subsequent compositional analyses focused on biofilms cultivated on glass only.

**Fig. 4.**
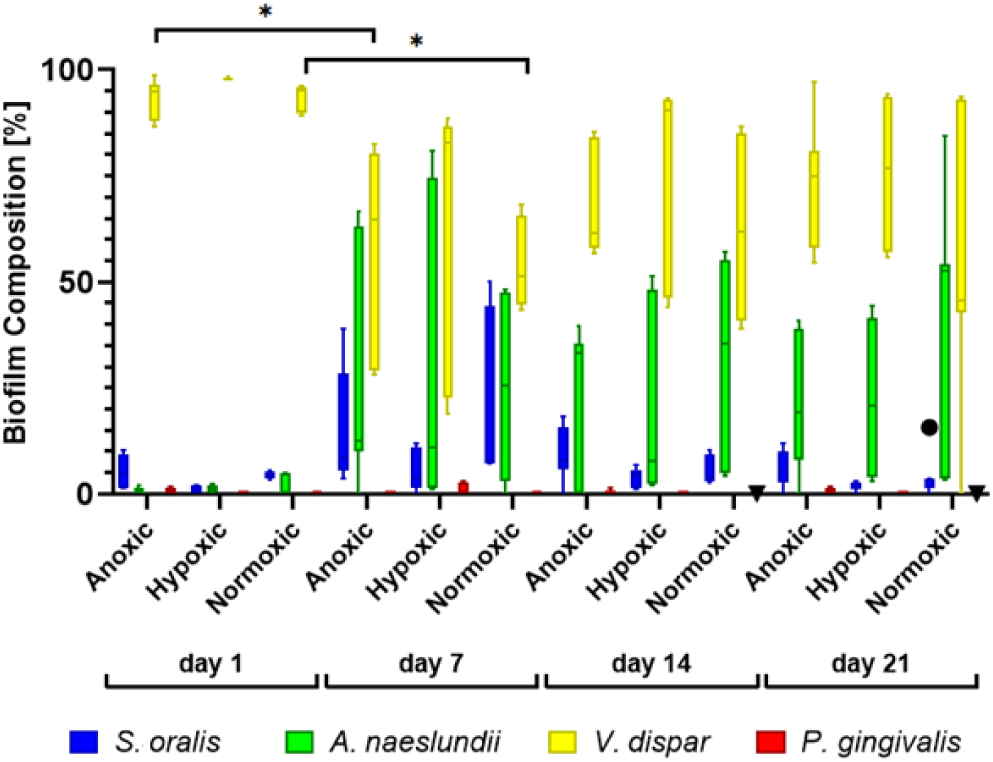
Tukey box plots showing the relative species composition of four-species biofilms cultivated under different oxygen conditions over 21 days. Biofilms were grown under anoxic (0%), hypoxic (1%), and normoxic (21%) conditions. * indicates statistically significant differences with *p* ≤ 0.05.

At day 1, biofilms were strongly dominated by *V. dispar*, accounting for nearly 100% of the detected bacterial population under all oxygen conditions. In contrast, *A. naeslundii* and *S. oralis* were present in low proportions (less than 5%), while *P. gingivalis* remained near or below the detection limit at this early time point. By day 7, changes in relative species proportions were observed under anoxic and normoxic conditions, whereas greater variability was detected under hypoxic conditions. Under both conditions, the relative abundance of *V. dispar* decreased significantly compared to day 1 to approx. 60% (p = 0.0163 and p = 0.0167, respectively), although it remained the most abundant species across all oxygen conditions at day 7. Concurrently, modest increases in *A. naeslundii* and *S. oralis* to 20-30% were observed under all cultivation conditions. These compositional patterns persisted through day 14 and day 21.

Overall, throughout the entire 21-day cultivation period and independently of the oxygen condition, the biofilm was dominated by *V. dispar*. *A. naeslundii* consistently represented the second most abundant species, and *S. oralis* maintained a low-to-moderate relative abundance, with slightly higher proportions observed under normoxic conditions at day 7. *P. gingivalis* remained a minor component under all conditions and time points examined.

### Distinct vertical and horizontal spatial species organization

Spatial species distribution within the biofilms was additionally visualized using Fluorescence *In Situ* Hybridization (FISH) combined with CLSM. The 3D reconstructions of z-stacks using the Imaris software allowed the assessment across different oxygen conditions, time points and vertical planes within the biofilm. Representative FISH images are shown in Fig. 5.

**Fig. 5.**
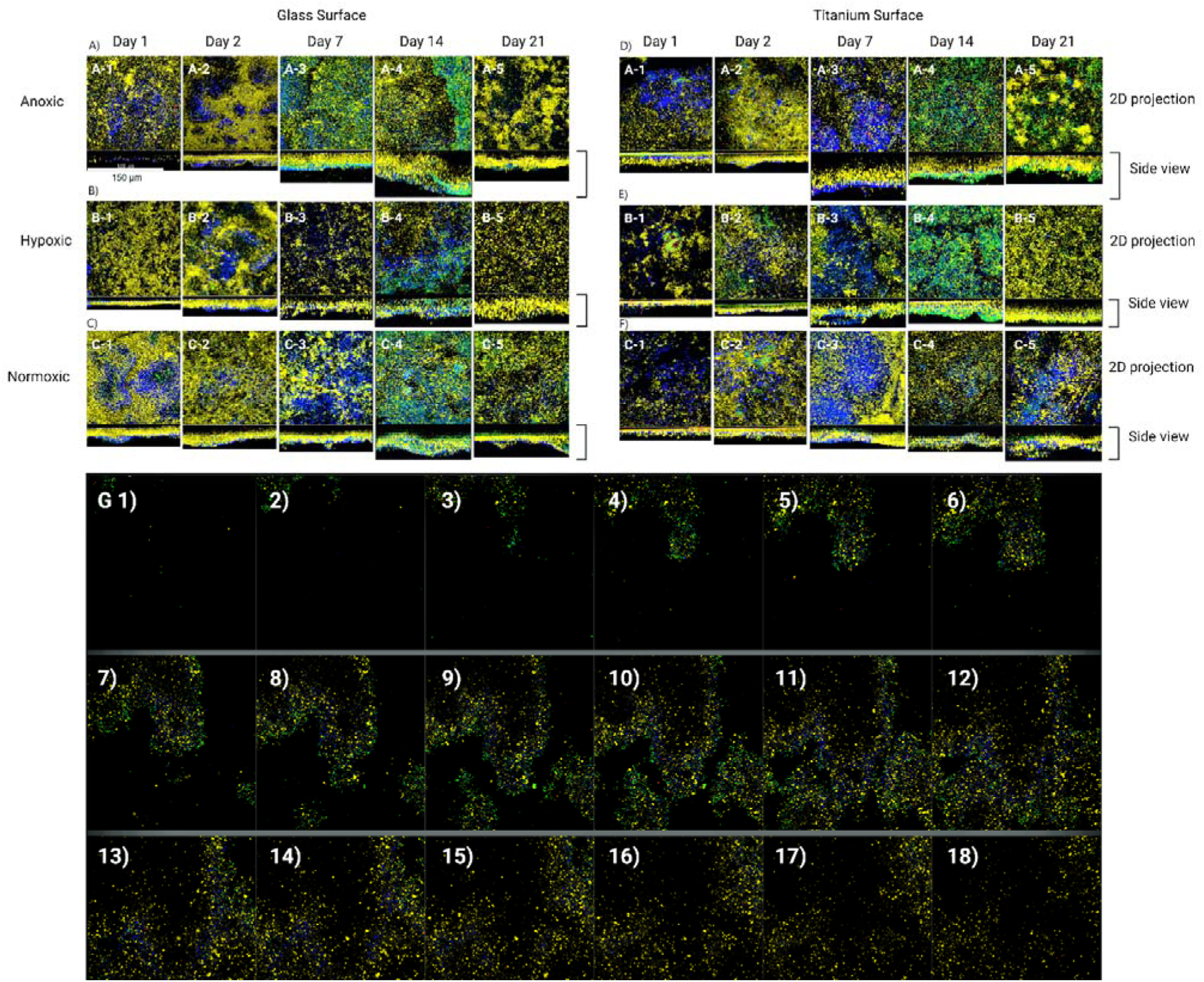
Representative microscopy images of spatial distribution of individual bacterial species within multispecies biofilms visualized by Fluorescence *In Situ* Hybridization (FISH). Blue = *S. oralis*, green = *A. naeslundii*, yellow = *V. dispar*, red = *P. gingivalis*. Biofilms were cultivated for 1, 2, 7, 14 and 21 days on on A) - C) glass or D) - F) titanium surfaces under A) D) anoxic, B) E) hypoxic, C) F) normoxic conditions. G) Representative images taken sorted by depth from bottom to top of biofilms cultivated under anoxic condition on glass for 14 days, showing the three-dimensional distribution of bacterial species (18 optical sections, z-step size = 2 µm). Images were acquired by confocal laser scanning microscopy (CLSM). Scale bar = 150 µm.

Across all conditions and time points examined, biofilms displayed a heterogeneous three-dimensional architecture, with species-specific spatial organization rather than uniform mixing. On glass surfaces (Fig. 5A–C), early-stage biofilms (day 1–2) were characterized by dispersed bacterial aggregates interspersed with small microcolonies. *V. dispar* signals (yellow) were broadly distributed throughout the biofilm, whereas *S. oralis* (blue) were observed in specific layers, and *A. naeslundii* (green) were observed in localized clusters. At middle and late cultivation stages (day 7–21), biofilms became thicker, showing an increase in overall biomass and more pronounced stratification along the vertical axis. On titanium surfaces (Fig. 5D–F), similar spatial patterns were observed across oxygen conditions. Early biofilms exhibited patchy bacterial coverage, followed by the development of thicker biofilm structures at later time points. As on glass surfaces, *V. dispar* signals were widely distributed within the biofilm matrix, whereas *S. oralis* and *A. naeslundii* showed a more localized organization. No pronounced qualitative differences in spatial organization between anoxic, hypoxic, and normoxic conditions were apparent by visual inspection. Across both surface types and all oxygen conditions, *P. gingivalis* (red) signals were mainly detected as isolated cells or small clusters throughout the cultivation period, co-aggregated with *V. dispar*.

The FISH-stained biofilm cultivated under anoxic condition on glass for 14 days (Fig. 5G) revealed distinct vertical stratification within the biofilm structure in mature biofilms. In a thick biofilm, *S. oralis* (blue) and *A. naeslundii* (green) were predominantly located in the upper biofilm layers, while *V. dispar* (yellow) and *P. gingivalis* (red) were primarily detected in the lower biofilm regions.

Overall, FISH analysis demonstrated reproducible species-specific spatial organization, including distinct horizontal and vertical distribution patterns that were independent of the surrounding oxygen conditions.

## Discussion

The onset of peri-implant mucositis and peri-implantitis is associated with changes in the local oxygen conditions. However, how these conditions influence the polymicrobial biofilm that drives these infections remains poorly understood. The present study analysed a defined oral multispecies biofilm model and demonstrates its structural and compositional resilience to varying oxygen conditions. While external oxygen availability influenced early biofilm volume and membrane integrity, the community consistently maintained a *V. dispar*-dominated structure across all oxygen conditions. In parallel, pathogenic and oxygen-sensitive *P. gingivalis* also persisted in low proportions. This suggests that once the internal framework of the community is established, it becomes compositionally robust against environmental oxygen fluctuations and by this potentially paves the way for infection development.

Within the present study, three different oxygen conditions were analyzed. For normoxic conditions, 21% oxygen were applied, as found in oral cavity. In contrast, hypoxic conditions contained 1% to simulate peri-implant niches. Finally, anoxic conditions represented severe infection sites. Regarding biofilm morphology and viability, normoxic conditions promoted higher biofilm volume during the first two days of cultivation and supported higher membrane integrity throughout the experimental period compared to hypoxic and anoxic conditions. These effects were most significant during early biofilm development and diminished as the biofilms matured, with volume differences largely disappearing after day 7. In contrast, hypoxic and anoxic conditions resulted in comparable biofilm volumes and membrane integrity profiles, indicating that intermediate oxygen availability difference did not lead to a distinct difference for biofilm growth and physiology in this system.

Initially, the observation that normoxic conditions supported higher membrane integrity and early biofilm growth appears counterintuitive, given that the model consists exclusively of facultative or obligate anaerobic species. However, this phenomenon is consistent with literatures describing that not all “strictly anaerobic” bacteria respond uniformly to oxygen exposure (Uesugi, 1978) (Gerritse, 1992) (Wicaksono, 2020). While our study did not directly characterize specific metabolic pathways, previous research has indicated that *Veillonella* species can neutralize reactive oxygen species (ROS) from early colonizers through catalase or peroxidase activities, effectively lowering the local redox potential (Zhou P. M., 2021). In such cases, oxygen exposure does not necessarily inhibit growth but may transiently activate stress responses or metabolic adaptations that facilitate biofilm community establishment. The sustained physiological robustness observed in our normoxic biofilms is congruent with the potential formation of such protective microenvironments. In the context of a developing biofilm, early exposure to oxygen may accelerate the formation of protective microenvironments as kind of stress reaction by stimulating metabolic activity, matrix production, or cooperative interactions among community members (Guggenheim, 2001) (Chalmers, 2008).

Importantly, these oxygen-dependent effects on growth and viability were not accompanied by major shifts in species composition as assessed by qRT-PCR. Across all oxygen conditions and time points, *V. dispar* remained the dominant species, whereas *A. naeslundii* and *S. oralis* increased modestly during middle and later stages. Notably, *P. gingivalis* persisted as a minor but viable community member (approximately 1%) across all oxygen gradients. This compositional resilience is likely reinforced by the vertical stratification observed via FISH, with *S. oralis* and *A. naeslundii* preferentially localized in upper biofilm layers, while *V. dispar* and *P. gingivalis* were more localized in deeper regions. This vertical stratification aligns with classical ecological models of oral biofilms, in which aerotolerant facultative anaerobic species near the surface support the formation of oxygen-reduced microenvironments in deeper layers (Loesche, 1983) (Takahashi, 2015). Such spatial structuring provides a mechanistic framework for understanding why external oxygen availability had limited impact on community composition. Even under normoxic conditions, this oxygen gradient produced by microbiomes is likely to develop rapidly within the biofilm, rendering deeper regions effectively anaerobic. The observed colocalization of *V. dispar* and *P. gingivalis* is consistent with previous studies suggesting that *Veillonella* species can provide nutritional support and serve as physical attachment partners for *P. gingivalis* (Zhou P. L., 2015). From a clinical standpoint, the persistence of *P. gingivalis* under normoxic conditions in our model is a critical finding: it suggests that a *V. dispar*-dominated matrix acts as a sanctuary for pathogens. Our results suggest that this interspecies coupling provides a sufficiently sheltered niche to ensure the long-term persistence of oxygen-sensitive pathogens, regardless of external oxygen tension.

A key feature of this biofilm model is the persistent dominance of *V. dispar*, which differs from previously reported *Streptococcus*-dominant models (Kommerein, 2017) (Heine, 2025), irrespective of oxygen conditions. By employing a standardized 24 h liquid pre-culture following plate recovery, we lowered the metabolic activity of *S. oralis* (Fig. S-1) and established a community equilibrium centered on *V. dispar*. Given the strong growth phase dependent metabolic reprogramming of *V. dispar* and the physiological state of the inoculum (Zhang, 2023), carryover effects from preculture are likely to result in a disproportionate influence during the initial stages of community assembly; however, the exact nature of these alterations requires further experimental validation. Once this *Veillonella*-centric community is established, sustained selective pressures, which ranged from anoxia to atmospheric oxygen, failed to override its dominance. This could be deeply rooted in the metabolic interdependencies among member species (Jakubovics, 2015). Within this syntrophic framework, *Actinomyces* and *Streptococcus* species initially colonize and assist the integration of *Veillonella* through specific coaggregation interactions (Hughes, 1988). *Streptococcus* generates lactate as a fermentation by-product, which serves as the preferred substrate for *Veillonella*. The subsequent sequestration of lactate mitigates end-product inhibition, thereby enhancing the metabolic activity of the streptococcal population (Chalmers, 2008) (Gross, 2012) (Mashima, 2015). Furthermore, recent work has demonstrated that *V. dispar* exhibits strong metabolic plasticity across growth phases, actively reprogramming its lactate metabolism and short-chain fatty acid production during transitions from exponential to stationary phase (Zhang, 2023). Beyond serving as a metabolic hub (Kolenbrander, 2006) (Persson, 2014), most *Veillonella* species utilize catalase to neutralize reactive oxygen species (ROS) produced by early colonizers. By detoxifying ROS and effectively lowering the local redox potential, *Veillonella* species establish a hypoxic niche (Zhou P. L., 2017). This capacity for adaptive metabolic rerouting and ROS detoxification allows *V. dispar* to actively stabilize the biofilm environment, recruit oxygen-sensitive pathogens like *P. gingivalis*, and function as a robust biological buffer against varying oxygen stresses. The clinical implication of this “biological buffering” is significant for therapeutic strategies, as it allows pathogenic members to maintain or expand their proportion regardless of external oxygen tensions. This underscores the decisive “window of opportunity” in clinical practice, specifically the first few days following implant placement or professional cleaning. Nevertheless, further investigations incorporating different inoculum preconditioning protocols are required to systematically evaluate the influence of initial cultivation conditions on biofilm development.

In conclusion, external oxygen availability modulates early physiological parameters such as biofilm volume and membrane integrity, but does not override the structural and compositional framework established during initial community assembly. In this four-species model, the metabolic resilience of *V. dispar* and the resulting internal microenvironments may emerge as the primary drivers of biofilm stability. These results suggest that clinical oxygen levels alone may not dictate the presence of pathogens and highlight the need to target the metabolic communications of the biofilm to overcome its environmental resilience.

Building on the observed structural and compositional resilience of this *V. dispar*-dominant model, future investigations should systematically compare its developmental dynamics with traditional *S. oralis*-dominated biofilm. In addition, applying multi-omics approaches such as transcriptomics, proteomics, and metabolomics will be essential to elucidate the underlying metabolic pathways, particularly the differential expression of oxidative stress response genes. Moreover, incorporating additional bridging species and host-derived factors, including inflammatory mediators and immune cells, will help disentangle the complex interplay between initial community physiology and environmental pressures. Such integrated approaches will enhance the translational value of *in vitro* oral multispecies biofilm model, providing a deeper understanding of the persistent nature of microbial consortia in peri-implant infections.

## Acknowledgments

Incubation of biofilm under hypoxic conditions was performed by using a Whitley H35 HEPA hypoxystation at Institute for Medical Microbiology and Hospital Epidemiology, Hannover Medical School and we would like to thank Maria Hänel for her support and advice. We would also like to thank Diana Strauch and Rainer Schreeb for their technical assistance.

## Data availability statement

The data that support the findings of this study are available from the corresponding author upon reasonable request.

## Declaration of Generative AI Use

Gemini 3.1 Pro was used during the drafting of this manuscript exclusively to improve spelling, grammar, and linguistic flow. The AI tool was not used to generate data, analyze results, or draw scientific conclusions. The author(s) reviewed all modifications and assume(s) full responsibility for the accuracy and originality of the published article.

## Supplementary data

**Fig. S-1.**
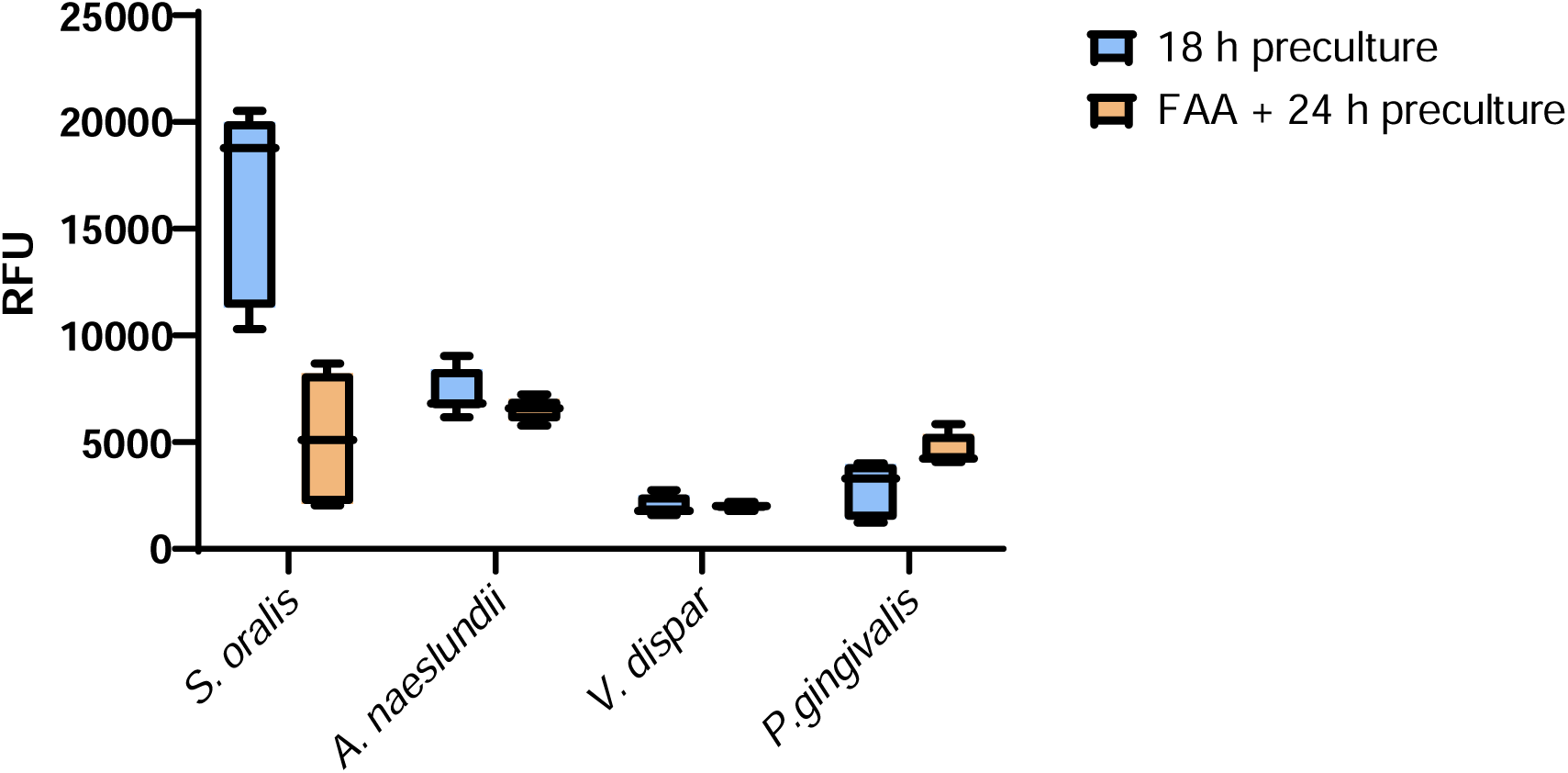
Biofilm metabolic activity of *S. oralis*, *A. naeslundii*, *V. dispar*, and *P. gingivalis* after 18 h BHI preculture (blue) and 3 days on FAA plates + 24 h BHI preculture (orange) tested with resazurin.

**Table S-1:**
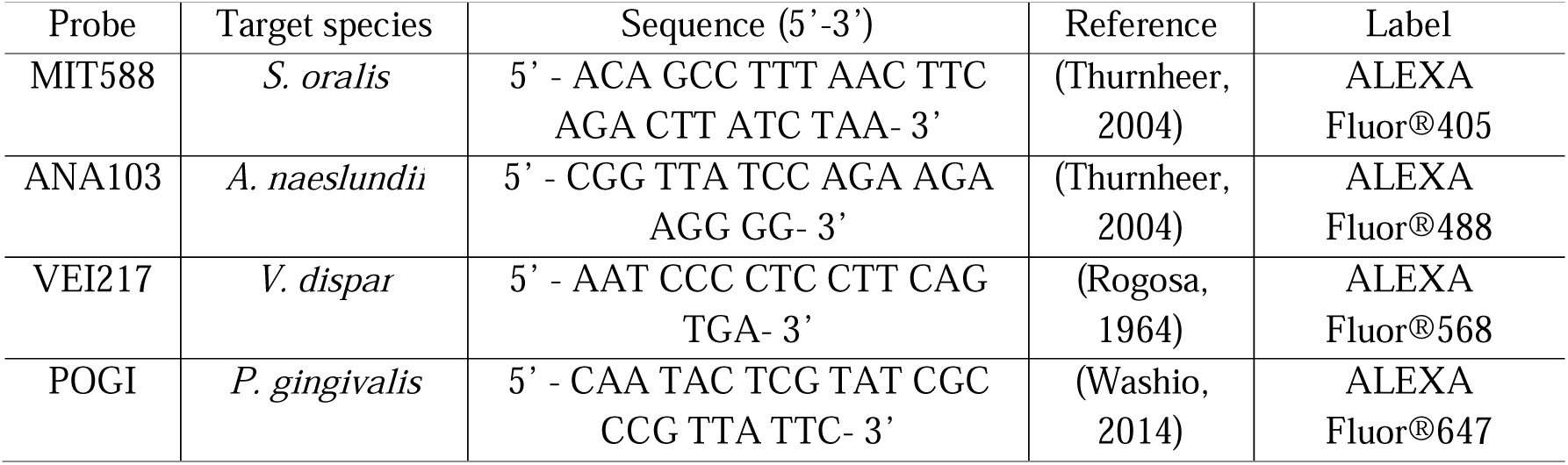
Species-specific 16S rRNA probes for fluorescence *in situ* hybridization.

**Table S-2:**
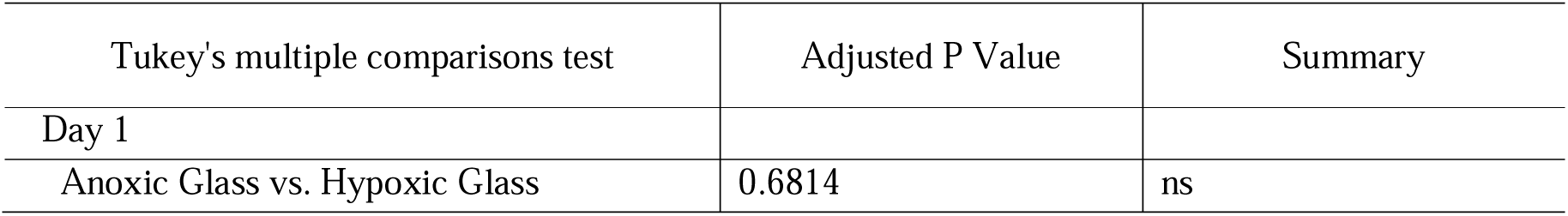
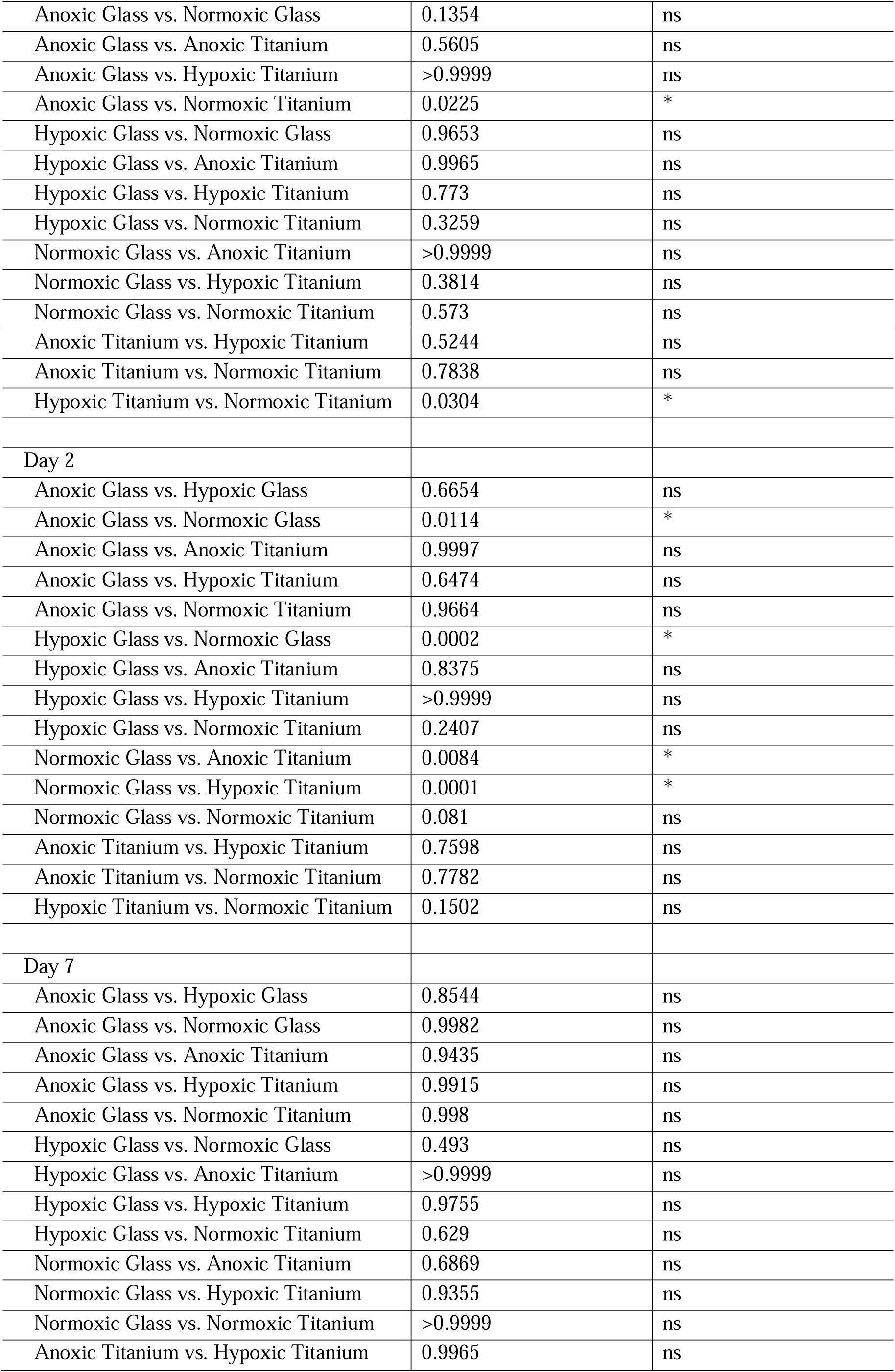
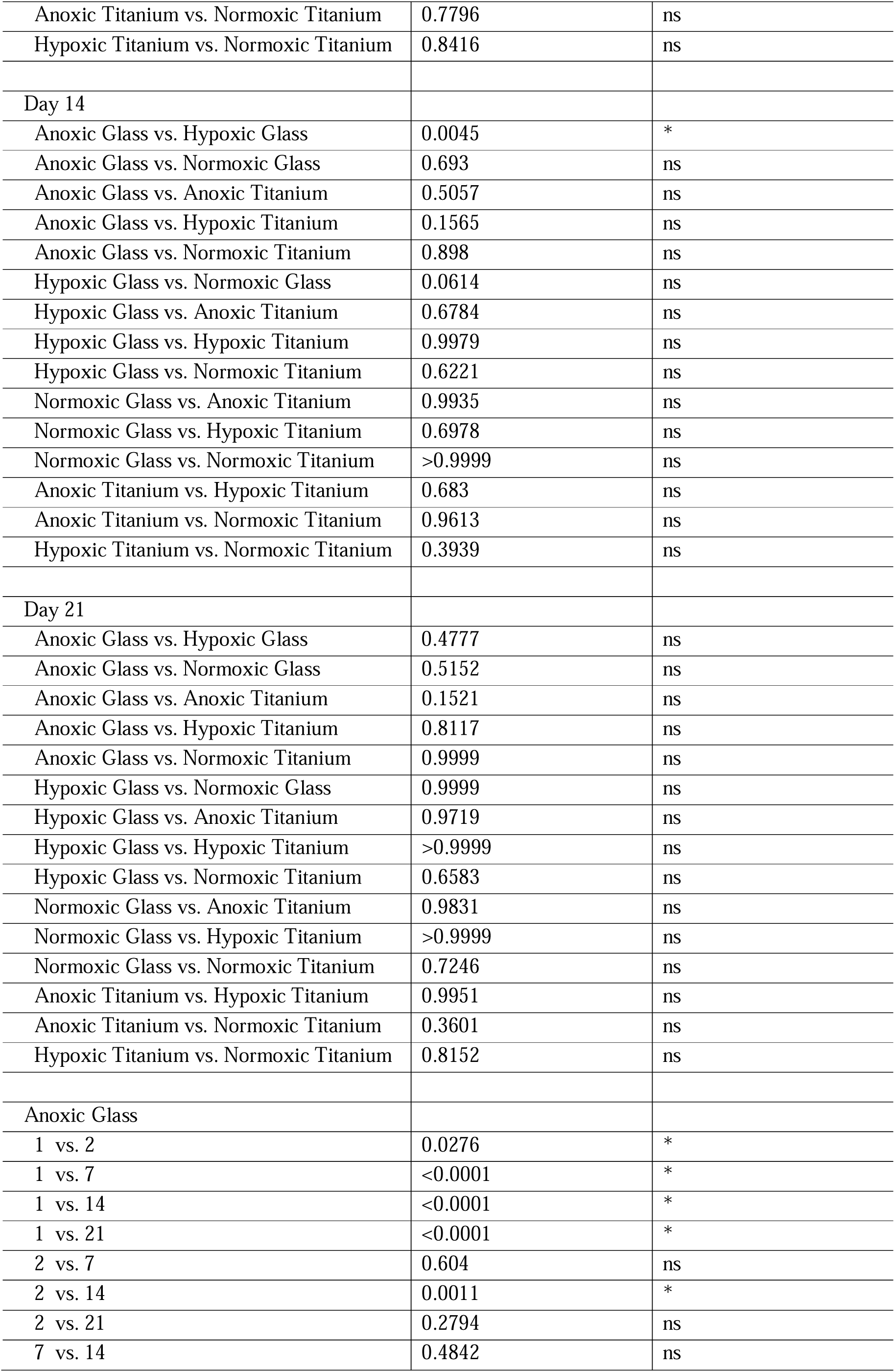
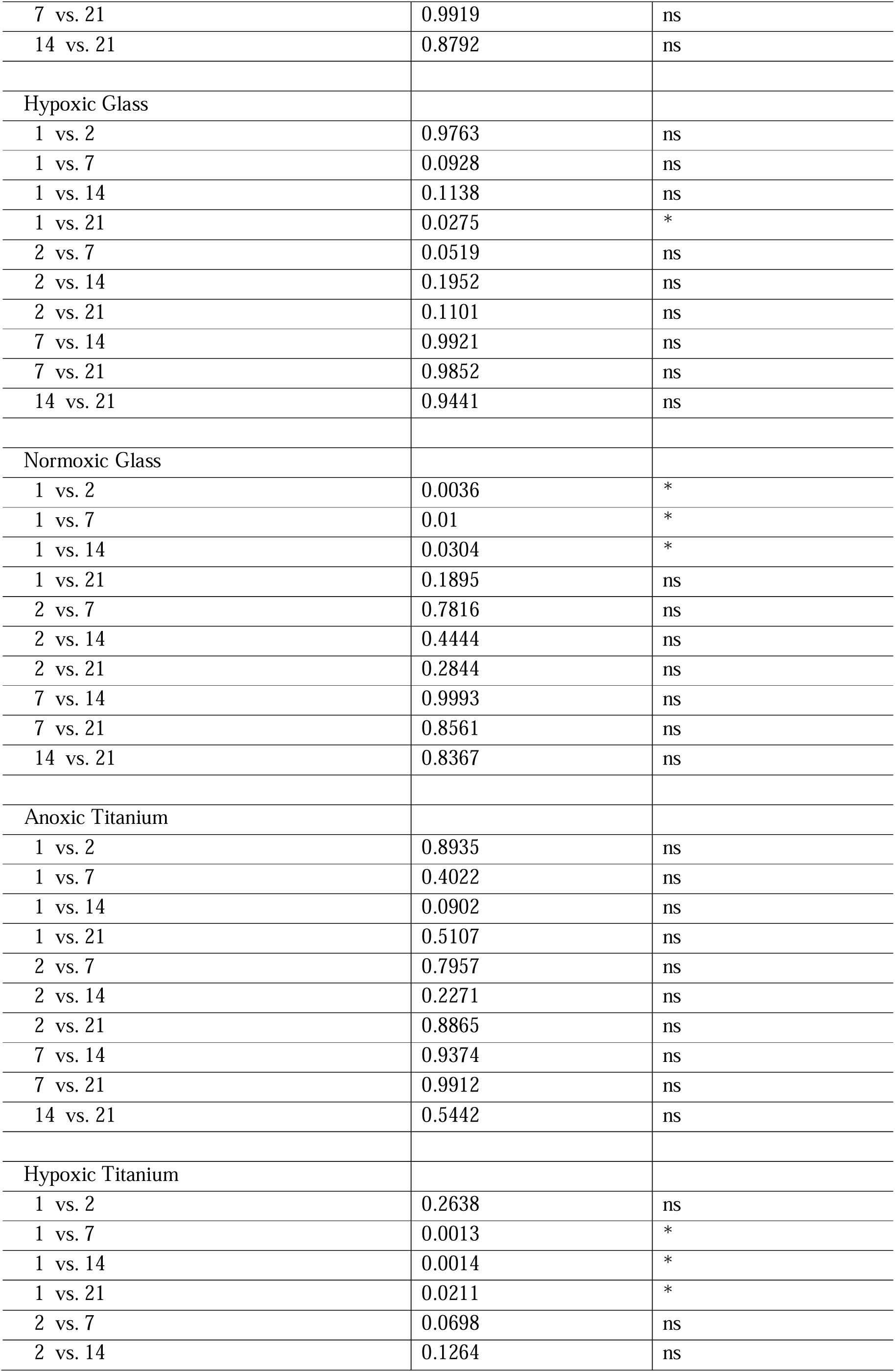
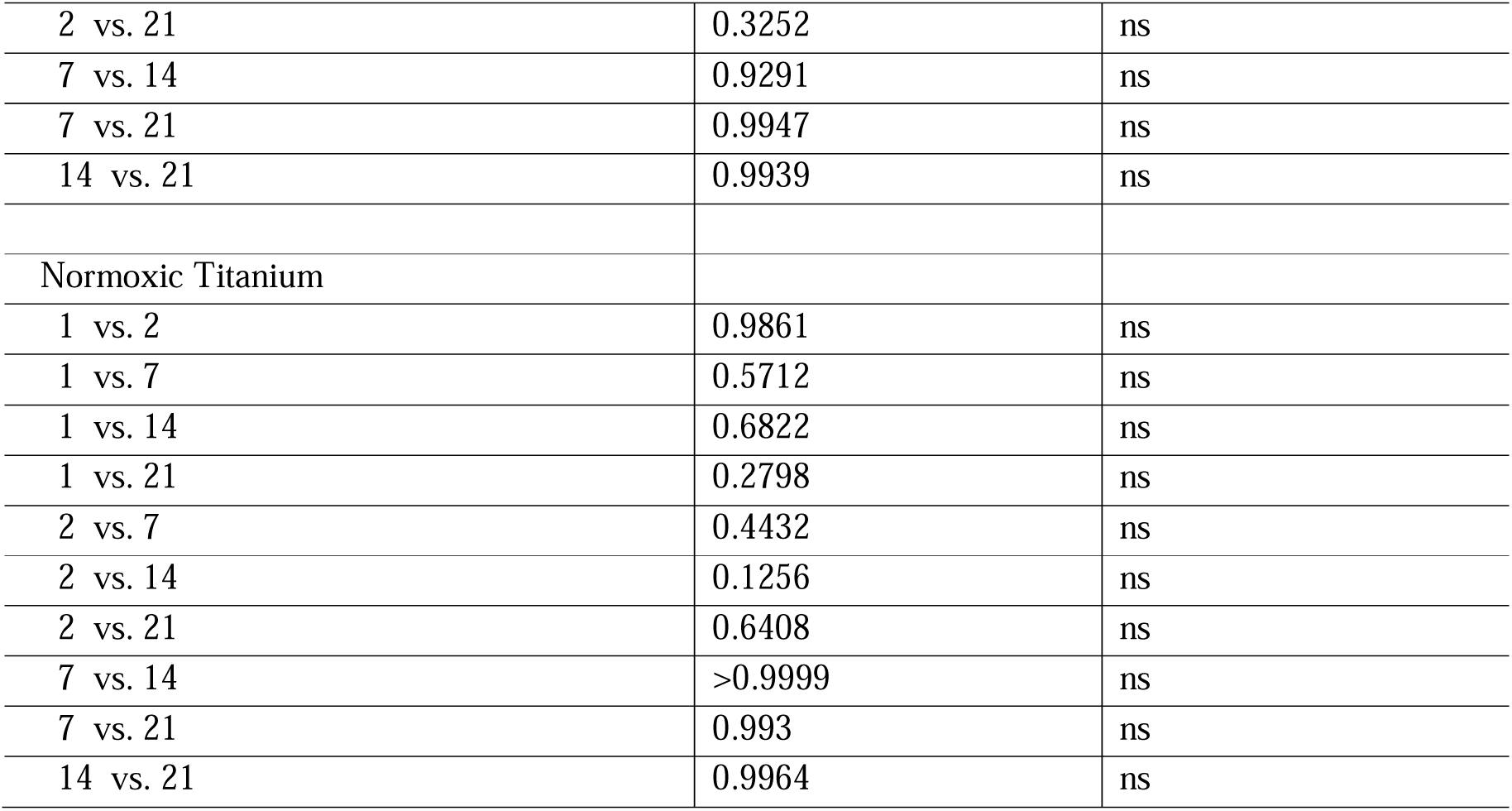
2way ANOVA comparisons of biofilm volume between different oxygen conditions, followed by a Tukey’s post-hoc test for multiple comparisons and p-value correction, with statistical significance defined at α = 0.05

**Table S-2:** 2way ANOVA comparisons of biofilm membrane integrity between different oxygen conditions, followed by a Tukey’s post-hoc test for multiple comparisons and p-value correction, with statistical significance defined at α = 0.05

| Tukey's multiple comparisons test | Adjusted P Value | Summary |
| --- | --- | --- |
| Day 1 |  |  |
| Anoxic Glass vs. Hypoxic Glass | 0.0092 | * |
| Anoxic Glass vs. Normoxic Glass | 0.1294 | ns |
| Anoxic Glass vs. Anoxic Titanium | 0.0058 | * |
| Anoxic Glass vs. Hypoxic Titanium | 0.0469 | * |
| Anoxic Glass vs. Normoxic Titanium | 0.0359 | * |
| Hypoxic Glass vs. Normoxic Glass | >0.9999 | ns |
| Hypoxic Glass vs. Anoxic Titanium | 0.9995 | ns |
| Hypoxic Glass vs. Hypoxic Titanium | >0.9999 | ns |
| Hypoxic Glass vs. Normoxic Titanium | 0.9902 | ns |
| Normoxic Glass vs. Anoxic Titanium | >0.9999 | ns |
| Normoxic Glass vs. Hypoxic Titanium | >0.9999 | ns |
| Normoxic Glass vs. Normoxic Titanium | 0.98 | ns |
| Anoxic Titanium vs. Hypoxic Titanium | >0.9999 | ns |
| Anoxic Titanium vs. Normoxic Titanium | 0.9888 | ns |
| Hypoxic Titanium vs. Normoxic Titanium | 0.9902 | ns |
| Day 2 |  |  |
| Anoxic Glass vs. Hypoxic Glass | 0.9992 | ns |
| Anoxic Glass vs. Normoxic Glass | 0.0003 | * |
| Anoxic Glass vs. Anoxic Titanium | 0.89 | ns |
| Anoxic Glass vs. Hypoxic Titanium | >0.9999 | ns |
| Anoxic Glass vs. Normoxic Titanium | 0.0014 | * |
| Hypoxic Glass vs. Normoxic Glass | 0.0006 | * |
| Hypoxic Glass vs. Anoxic Titanium | 0.94 | ns |
| Hypoxic Glass vs. Hypoxic Titanium | 0.9713 | ns |
| Hypoxic Glass vs. Normoxic Titanium | 0.0094 | * |
| Normoxic Glass vs. Anoxic Titanium | 0.0001 | * |
| Normoxic Glass vs. Hypoxic Titanium | <0.0001 | * |
| Normoxic Glass vs. Normoxic Titanium | 0.6792 | ns |
| Anoxic Titanium vs. Hypoxic Titanium | 0.5737 | ns |
| Anoxic Titanium vs. Normoxic Titanium | 0.0042 | * |
| Hypoxic Titanium vs. Normoxic Titanium | 0.0008 | * |
| Day 7 |  |  |
| Anoxic Glass vs. Hypoxic Glass | 0.9972 | ns |
| Anoxic Glass vs. Normoxic Glass | 0.1309 | ns |
| Anoxic Glass vs. Anoxic Titanium | 0.6797 | ns |
| Anoxic Glass vs. Hypoxic Titanium | 0.6736 | ns |
| Anoxic Glass vs. Normoxic Titanium | 0.0093 | * |
| Hypoxic Glass vs. Normoxic Glass | 0.0403 | * |
| Hypoxic Glass vs. Anoxic Titanium | 0.831 | ns |
| Hypoxic Glass vs. Hypoxic Titanium | 0.6204 | ns |
| Hypoxic Glass vs. Normoxic Titanium | 0.019 | * |
| Normoxic Glass vs. Anoxic Titanium | 0.8552 | ns |
| Normoxic Glass vs. Hypoxic Titanium | 0.9441 | ns |
| Normoxic Glass vs. Normoxic Titanium | 0.5785 | ns |
| Anoxic Titanium vs. Hypoxic Titanium | >0.9999 | ns |
| Anoxic Titanium vs. Normoxic Titanium | 0.2428 | ns |
| Hypoxic Titanium vs. Normoxic Titanium | 0.422 | ns |
| Day 14 |  |  |
| Anoxic Glass vs. Hypoxic Glass | 0.4911 | ns |
| Anoxic Glass vs. Normoxic Glass | <0.0001 | * |
| Anoxic Glass vs. Anoxic Titanium | 0.2734 | ns |
| Anoxic Glass vs. Hypoxic Titanium | 0.0221 | * |
| Anoxic Glass vs. Normoxic Titanium | 0.025 | * |
| Hypoxic Glass vs. Normoxic Glass | <0.0001 | * |
| Hypoxic Glass vs. Anoxic Titanium | 0.9647 | ns |
| Hypoxic Glass vs. Hypoxic Titanium | 0.354 | ns |
| Hypoxic Glass vs. Normoxic Titanium | 0.3354 | ns |
| Normoxic Glass vs. Anoxic Titanium | 0.0869 | ns |
| Normoxic Glass vs. Hypoxic Titanium | 0.5019 | ns |
| Normoxic Glass vs. Normoxic Titanium | 0.3038 | ns |
| Anoxic Titanium vs. Hypoxic Titanium | 0.7769 | ns |
| Anoxic Titanium vs. Normoxic Titanium | 0.9847 | ns |
| Hypoxic Titanium vs. Normoxic Titanium | 0.9988 | ns |
| Day 21 |  |  |
| Anoxic Glass vs. Hypoxic Glass | 0.9938 | ns |
| Anoxic Glass vs. Normoxic Glass | 0.0219 | * |
| Anoxic Glass vs. Anoxic Titanium | 0.9982 | ns |
| Anoxic Glass vs. Hypoxic Titanium | 0.9997 | ns |
| Anoxic Glass vs. Normoxic Titanium | 0.4769 | ns |
| Hypoxic Glass vs. Normoxic Glass | 0.0091 | * |
| Hypoxic Glass vs. Anoxic Titanium | 0.8792 | ns |
| Hypoxic Glass vs. Hypoxic Titanium | >0.9999 | ns |
| Hypoxic Glass vs. Normoxic Titanium | 0.2194 | ns |
| Normoxic Glass vs. Anoxic Titanium | 0.004 | * |
| Normoxic Glass vs. Hypoxic Titanium | 0.0537 | ns |
| Normoxic Glass vs. Normoxic Titanium | 0.7273 | ns |
| Anoxic Titanium vs. Hypoxic Titanium | 0.9712 | ns |
| Anoxic Titanium vs. Normoxic Titanium | 0.1187 | ns |
| Hypoxic Titanium vs. Normoxic Titanium | 0.6064 | ns |
| Anoxic Glass |  |  |
| 1 vs. 2 | 0.0041 | * |
| 1 vs. 7 | 0.0151 | * |
| 1 vs. 14 | 0.0182 | * |
| 1 vs. 21 | 0.0969 | ns |
| 2 vs. 7 | >0.9999 | ns |
| 2 vs. 14 | 0.9277 | ns |
| 2 vs. 21 | 0.434 | ns |
| 7 vs. 14 | 0.8286 | ns |
| 7 vs. 21 | 0.3232 | ns |
| 14 vs. 21 | 0.4727 | ns |
| Hypoxic Glass |  |  |
| 1 vs. 2 | 0.9359 | ns |
| 1 vs. 7 | 0.7344 | ns |
| 1 vs. 14 | 0.9806 | ns |
| 1 vs. 21 | 0.9406 | ns |
| 2 vs. 7 | 0.9593 | ns |
| 2 vs. 14 | 0.8504 | ns |
| 2 vs. 21 | 0.734 | ns |
| 7 vs. 14 | 0.0208 | * |
| 7 vs. 21 | 0.0676 | ns |
| 14 vs. 21 | >0.9999 | ns |
| Normoxic Glass |  |  |
| 1 vs. 2 | 0.0005 | * |
| 1 vs. 7 | 0.7869 | ns |
| 1 vs. 14 | 0.0032 | * |
| 1 vs. 21 | 0.0164 | * |
| 2 vs. 7 | 0.0013 | * |
| 2 vs. 14 | 0.7996 | ns |
| 2 vs. 21 | 0.4121 | ns |
| 7 vs. 14 | 0.0103 | * |
| 7 vs. 21 | 0.0474 | * |
| 14 vs. 21 | 0.9681 | ns |
| Anoxic Titanium |  |  |
| 1 vs. 2 | 0.9986 | ns |
| 1 vs. 7 | 0.9996 | ns |
| 1 vs. 14 | 0.3749 | ns |
| 1 vs. 21 | 0.9989 | ns |
| 2 vs. 7 | >0.9999 | ns |
| 2 vs. 14 | 0.6208 | ns |
| 2 vs. 21 | >0.9999 | ns |
| 7 vs. 14 | 0.5354 | ns |
| 7 vs. 21 | >0.9999 | ns |
| 14 vs. 21 | 0.5528 | ns |
| Hypoxic Titanium |  |  |
| 1 vs. 2 | 0.8593 | ns |
| 1 vs. 7 | 0.9719 | ns |
| 1 vs. 14 | 0.0005 | * |
| 1 vs. 21 | 0.8705 | ns |
| 2 vs. 7 | 0.8503 | ns |
| 2 vs. 14 | 0.0195 | * |
| 2 vs. 21 | 0.4324 | ns |
| 7 vs. 14 | 0.095 | ns |
| 7 vs. 21 | 0.9893 | ns |
| 14 vs. 21 | 0.3967 | ns |
| Normoxic Titanium |  |  |
| 1 vs. 2 | 0.0087 | * |
| 1 vs. 7 | 0.1992 | ns |
| 1 vs. 14 | 0.0806 | ns |
| 1 vs. 21 | 0.0493 | * |
| 2 vs. 7 | 0.8621 | ns |
| 2 vs. 14 | 0.6021 | ns |
| 2 vs. 21 | 0.5019 | ns |
| 7 vs. 14 | 0.9873 | ns |
| 7 vs. 21 | 0.9938 | ns |
| 14 vs. 21 | >0.9999 | ns |

